# Insights Into The Early Diversification of *bla*_NDM-1_ Genetic Contexts Following its Emergence in *Acinetobacter* species and Spread to Enterobacterales

**DOI:** 10.64898/2026.10.22.760963

**Authors:** Nancy Allard, Dalila Touati, Nathan Faille, Frédéric Grenier, Emy Beauchemin-Lauzon, Simon Lévesque, Sébastien Rodrigue, Louis-Patrick Haraoui

## Abstract

Since its emergence, the New Delhi metallo-β-lactamase (NDM) has become a major global antimicrobial resistance threat, enabling bacterial hosts to resist most β-lactam antibiotics, including carbapenems, and spreading widely across bacterial species and geographic regions. Believed to have originated in *Acinetobacter* spp., *bla*_NDM_ subsequently spread to Enterobacterales through mobile genetic elements, despite barriers to lateral gene transfer between these phylogenetically distant taxa. To investigate the genetic diversity and mechanisms underlying this interspecies transfer and subsequent global expansion, we sequenced and characterized 13 *bla*_NDM-1_-positive Gram-negative bacterial isolates collected in India in 2007, among the earliest known isolates harboring *bla*_NDM-1_. Clonal diversity was observed among both *Escherichia coli* and *Klebsiella pneumoniae* isolates. Comprehensive genomic analysis revealed that *bla*_NDM-1_ was consistently embedded within a conserved multidrug resistance region, often associated with mobile genetic elements involved in lateral gene transfer. Notably, the insertion sequence IS*Aba14* was identified in 6 of the 13 sequences, suggesting a potential role in the early dissemination of *bla*_NDM-1_ among diverse bacterial hosts. These findings contribute to our understanding of the evolutionary origins and early genetic context of *bla*_NDM-1_, highlighting the importance of continued genomic surveillance and functional studies to better understand the persistence and dissemination of carbapenem resistance.

**Impact Statement:** This study characterizes 13 previously undescribed genetic contexts of the New Delhi metallo-β-lactamase-1 gene (*bla*_NDM-1_) in Enterobacterales isolates collected in India in 2007. These isolates represent some of the earliest known *bla*_NDM-1_-positive Enterobacterales and are the earliest collected *bla*_NDM-1_-positive Enterobacterales for which whole-genome sequences are now available. Their genetic contexts include diverse combinations of antibiotic resistance genes and associated mobile genetic elements. Notably, the insertion sequence IS*Aba14* was identified in 6 of the 13 sequences, highlighting its frequent association with *bla*_NDM-1_ during the early stages of its dissemination. Together, these findings reveal substantial genetic diversity surrounding *bla*_NDM-1_ early in its emergence and suggest that diverse arrangements of mobile genetic elements may have contributed to its early dissemination across bacterial hosts and the subsequent spread of carbapenem resistance.

## Introduction

The use of antibiotics has revolutionized medicine and saved countless lives; however, their overuse and misuse have accelerated the emergence of antibiotic-resistant bacteria (1). <u>A</u>nti<u>m</u>icrobial resistance (AMR) has become a major global public health issue, complicating infection treatment and increasing the risks of disease spread, severe illness, and death (2). In bacteria, AMR can be acquired either vertically, through inherited mutations, or laterally via <u>m</u>obile <u>g</u>enetic <u>e</u>lements (MGEs) such as conjugative plasmids, transposons, integrons and insertion sequences (1,3). These MGEs often carry one or multiple <u>a</u>ntimicrobial resistance <u>g</u>enes (ARGs), playing a pivotal role in spreading resistance both within and across bacterial species (4). A striking example of an emerging ARG with worldwide distribution is the <u>N</u>ew <u>D</u>elhi <u>m</u>etallo-β-lactamase gene (*bla*_NDM-1_). This gene confers resistance to a broad range of β-lactam antibiotics, including penicillins, cephalosporins and carbapenems—a class often considered a last resort for treating severe bacterial infections (5). Since its first description in a patient epidemiologically linked to the Indian subcontinent, *bla*_NDM-1_ has disseminated globally and is now reported in more than 62 countries across all inhabited continents (6). Its worldwide spread among both healthcare-associated and community-associated bacteria has been strongly linked with international travel (7), including medical tourism, and the over-the-counter availability of broad-spectrum antibiotics in certain regions (8). Like other <u>m</u>etallo-<u>β</u>-lactamases (MBLs), *bla*_NDM-1_ is often found on MGEs and is typically located within highly recombining and transposon-rich regions, which also contributed to the spread of this resistance gene (7,9,10).

The *bla*_NDM-1_ gene was first reported in 2009 following its isolation in 2008 and subsequent characterization (11). Although newly discovered at that time, several unique features suggested that this gene had been evolving for decades, with recent estimates placing its origin before 1985 (10). Based on its distinctive biochemical properties and its low amino acid identity (32.4%) to the closest known β-lactamase, <u>V</u>erona integron-encoded <u>m</u>etallo-β-lactamase <u>2</u> (VIM-2), *bla*_NDM-1_ was recognized as the founding member of a new family designated New Delhi metallo-β-lactamase (11). It belongs to the Ambler class B metallo-β-lactamase that requires at least one zinc ion in its active site, thereby defining these enzymes as MBLs (12). The NDM-1 forms a lipoprotein anchored to the outer membrane of Gram-negative bacteria, unlike other MBLs (such as IMP and VIM), which are soluble periplasmic proteins (13). Clinically, NDM-1 confers resistance to nearly all β-lactams except aztreonam, posing a major challenge for treatment options (14).

Initially identified in a *Klebsiella pneumoniae* strain from a urinary tract infection, *bla*_NDM-1_ was later detected in an *Escherichia coli* strain from the feces of the same patient who had lived in Sweden for many years and frequently travelled to New Delhi, India (11,15). Another notable observation was that *bla*_NDM-1_ was found in distinct genetic contexts in the two bacterial species from which it was isolated (11). Since its initial description, *bla*_NDM-1_ has disseminated across phylogenetically diverse Gram-negative bacteria, predominantly Enterobacterales and *Acinetobacter* spp., with its presence documented in at least 11 bacterial families within the Gammaproteobacteria (16), including *Pseudomonas* spp., *Stenotrophomonas* spp., *Aeromonas* spp., and *Vibrio* (5,17). To date, 99 distinct variants of the NDM enzyme have been reported (http://www.bldb.eu/, accessed October 18, 2026), including approximately 40 identified within the past year. These variants exhibit minor amino acid substitutions that may affect enzymatic activity and alter resistance phenotypes (18).

Because of its prevalence among carbapenem-resistant Enterobacterales (19), *bla*_NDM-1_ has been the focus of extensive investigations into its origin, evolution and dissemination. Unlike many successful resistance determinants, the global spread of *bla*_NDM-1_ does not appear to be driven by the expansion of a single bacterial lineage, but rather repeated lateral gene transfer events involving diverse mobile genetic elements and bacterial hosts (9). Current evidence suggests that *bla*_NDM-1_ originated within an *Acinetobacter* background (19–21), where it was embedded within Tn*125*, a composite transposon consisting of two copies of IS*Aba125* flanking the resistance locus (10,21). The remarkable conservation of IS*Aba125* sequences, either intact or truncated, among *bla*_NDM-1_-positive isolates supports the hypothesis that Tn*125* was the ancestral gene vehicle responsible for the global dissemination of *bla*_NDM-1_ (10). Consistent with this hypothesis, the earliest known *bla*_NDM-1_-positive isolate identified to date is *Acinetobacter junii* Ajun-H1-3, recovered from a blood culture in Israel in 2004 (19), predating *bla*_NDM-1_-positive *Acinetobacter* spp. isolated in India in 2005 (22).

Despite considerable progress in reconstructing the evolutionary history of *bla*_NDM-1_, the early genetic events underlying its emergence in Enterobacterales remain poorly understood. In particular, the evolutionary processes and genetic intermediates involved in the transfer of *bla*_NDM-1_ from an *Acinetobacter* background into Enterobacterales remain unclear. Because Enterobacterales ultimately became the major reservoir responsible for the global dissemination of *bla*_NDM-1_, identifying the genetic contexts associated with its earliest occurrences in this taxonomic order may provide important insights into the emergence, adaptation, and subsequent spread of this clinically important resistance determinant.

In this study, we reconstructed the evolutionary and dissemination history of *bla*_NDM-1_ through comparative analyses of 13 newly sequenced genetic contexts from isolates collected in India in 2007. These isolates belong to a collection that includes some of the earliest known *bla*_NDM-1_-positive Enterobacterales, contemporaneous with the earliest reports of *bla*_NDM-1_-positive Enterobacterales recovered in India during 2006-2007 (23). By examining these early genetic contexts, we sought to gain insight into the events associated with the establishment of *bla*_NDM-1_ in Enterobacterales following its emergence in *Acinetobacter*. Among these, we identified six closely related structures carrying IS*Aba14* in association with IS*Aba125* and the *bla*_NDM-1_ locus. The conservation of these IS*Aba14*-containing contexts across distinct isolates suggests that they may represent previously unrecognized evolutionary intermediates in the emergence of the *bla*_NDM-1_ region. Together, our findings refine the current evolutionary model of the spread of *bla*_NDM-1_ and highlight a potentially important contribution of IS*Aba14*-associated genetic structures in the emergence and diversification of this globally distributed resistance determinant.

## Methodology

### Bacterial isolates

Thirty-six archived carbapenem-resistant Gram-negative bacterial isolates collected in India in 2007 were obtained from JMI Laboratories, representing nine bacterial taxa including *Klebsiella pneumoniae* (n=14), *Enterobacter cloacae* (n=6), *Acinetobacter baumannii* (n=5), *Escherichia coli* (n=3), *Pseudomonas aeruginosa* (n=2), *Ralstonia pickettii* (n=2), *Leclercia adecarboxyla* (n=2), *Elizabethkingae* sp. (n=1), and *Serratia rubidaea* (n=1). The isolates originated primarily from New Delhi (n=13) and Mumbai (n=9), with additional isolates from Pune (n=5), Kolkata (n=5), Cochin (n=2), Manipal (n=1), and Rajkot (n=1). Detailed information regarding isolate sources and species identification is provided in Supplementary Table S1. Carbapenemase genes were screened by polymerase chain reaction (PCR) as previously described by Mataseje *et al*. (24), including *bla*_NDM-1_, *<u>K</u>lebsiella <u>p</u>neumoniae* <u>c</u>arbapenemase (*bla*_KPC_), <u>im</u>i<u>p</u>enemase (*bla*_IMP_), *bla*_VIM_, <u>G</u>uiana-<u>e</u>xtended-<u>s</u>pectrum β-lactamase (*bla*_GES_), *<u>S</u>erratia <u>m</u>arcescens* <u>e</u>nzyme (*bla*_SME_), <u>Oxa</u>cillinase (*bla*_OXA-48-like_), and <u>n</u>on-<u>m</u>etallo-<u>c</u>arbapenemase/<u>imi</u>penemase-related (*bla*_NMC/IMI_)-like. Among the 36 isolates, 13 were positive for *bla*_NDM-1_ and were selected for whole-genome sequencing using both short-read (Illumina) and long-read (<u>O</u>xford <u>N</u>anopore <u>T</u>echnologies (ONT)) platforms.

### Bacterial strains and growth conditions

Bacterial strains used in this study are listed in Supplementary Table S1. Bacterial strains were grown at 37°C in Miller’s lysogeny <u>b</u>roth (LB) in an orbital shaker/incubator set at 225 rpm or on LB agar medium in a static incubator. Genomic DNA was extracted from bacterial isolates using the Quick-DNA Magbead Plus Kit (Zymo Research) and subjected to both short- and long-read sequencing. For short-read sequencing, DNA libraries were prepared as described by Grenier *et al.* (19). Libraries were sequenced on an Illumina NextSeq 500 platform using 40 <u>b</u>ase <u>p</u>air (bp) paired-end chemistry at the *Plateforme RNomics* (https://rnomics.med.usherbrooke.ca/) of the *Université de Sherbrooke*. For long-read sequencing, genomic DNA was treated with the NEBNext Ultra II End Repair/dA-Tailing Module (<u>N</u>ew <u>E</u>ngland <u>B</u>iolabs, NEB). Sample-specific barcodes from the Native Barcoding Kit 96 V14 (ONT) were ligated using the NEBNext Ultra II Ligation Module (NEB). Following purification with Ampure XP beads (Beckman Coulter), barcoded samples were pooled and sequencing adapters were ligated using the same Ligation Module (NEB). The final library was purified with Ampure XP beads (Beckman Coulter), and sequenced on an R10.4.1 MinION Flow Cell using a MinIon Mk1B device (ONT).

### Genome assembly and annotation

Illumina reads were quality filtered and trimmed using fastp v0.21.0 with the parameters --cut_right --cut_window_size 4 --cut_mean_quality 20 --length_required 30 --detect_adapter_for_pe (25). ONT reads were filtered using Filtlong v0.2.1 with the parameters --min_length 1000 --keep_percent 95 (https://github.com/rrwick/Filtlong). Hybrid genome assemblies were generated using Unicycler v0.4.9 (26) with trimmed Illumina short-reads and ONT long-reads. Assemblies for which the *bla*_NDM-1_-containing replicon could not be recovered as a complete circular sequence were reassembled using Trycycler v0.5.3 (27). Consensus assemblies were generated from twelve subsampled assemblies produced using Flye v2.9.1-b178 (28), Miniasm-Minipolish (29,30) (Minimap2 v2.24-r1122, Miniasm v0.3-r179 and Minipolish v0.1.3 and Raven v1.8.1) (31). Contigs shorter than 500 bp were excluded from downstream analyses. Genome annotation was performed using Prokka v1.14.5 (32) with Pfam and TIGRFAM databases. Genes located within the genetic environments surrounding *bla*_NDM-1_ were manually curated when necessary. Sequence data have been generated and will be deposited in GeneBank/Sequence Read Archive (SRA) upon manuscript submission or acceptance.

### Taxonomic identification and characterization of genetic contexts

Taxonomic assignment of all sequenced isolates was performed using Kraken2 v2.0.9-beta (25) on assembled genomes. Genetic environments surrounding *bla*_NDM-1_ were examined using annotated assemblies and manually inspected using SnapGene v8.2.0. When gene annotations were uncertain, nucleotide sequences were queried against the NCBI database using BLASTN for confirmation. Antimicrobial resistance genes were identified using ResFinder v4.1 (33) and the Resistance Gene Identifier (RGI) v6.0.4, which queries the <u>C</u>omprehensive <u>A</u>ntibiotic <u>R</u>esistance <u>D</u>atabase (CARD) (https://card.mcmaster.ca/). Plasmid incompatibility groups were determined using PlasmidFinder. (33). Insertion <u>s</u>equences (ISs) were initially identified using ISFinder online (https://isfinder.biotoul.fr/blast.php). Unless otherwise specified, default search parameters were employed. Only ISs sharing at least 95% nucleotide identity with reference sequences and extending over at least 100 bp were retained for downstream analyses. During manuscript revision, the original ISfinder server became unavailable; therefore, IS identification was subsequently performed using the ISfinder implementation available through TnCentral (https://tncentral.ncc.unesp.br/ISfinder/search.php). Genetic regions of interest were visualized using SnapGene v8.2.0 and Clinker v0.0.31. Final figures were manually refined using Inkscape.

### Antibiograms

Nine of the 13 *bla*_NDM-1_-positive isolates (renamed Escol1, Klepne1, Klepne2, Klepne3, Esccol3, Klepne4, Klepne5, Klepne6 and Enthor1) were subjected to antimicrobial susceptibility testing using Sensititre GNX3F panels according to the manufacturer’s instructions (Thermo Fisher Scientific). Susceptibility data for the remaining four isolates were generated previously by the strain provider using broth microdilution and were included for comparative purposes. A total of 21 antibiotics were tested, including amikacin, ampicillin/sulbactam, cefepime, cefoxitin, ceftazidime, ciprofloxacin, colistin, doripenem, doxycycline, gentamicin, imipenem, levofloxacin, meropenem, minocycline, piperacillin/tazobactam, tigecycline, tobramycin, TMP/SMX, aztreonam, polymyxin B, and ticarcillin/clavulanic acid (Supplementary Table S2). As no established interpretive breakpoints are available for tigecycline, except for *E. coli*, MICs are reported without susceptibility categorization.

## Results

### Isolate collection and genetic location of *bla*_NDM-1_

Thirty-six archived carbapenem-resistant Gram-negative isolates collected in India in 2007 were obtained from JMI Laboratories and screened for the presence of *bla*_NDM-1_. PCR screening identified *bla*_NDM-1_ in 13 isolates, including six *K. pneumoniae*, three *E. cloacae*, three *E. coli*, and one *A. baumannii*. These isolates originated from New Delhi (n=7), Mumbai (n=4), and Pune (n=2), and were recovered from bloodstream (n=5), respiratory tract (n=4), and skin and soft tissue (n=4) infections. The other resistance genes found were a combination of OXA-48 and KPC (n=7), a combination of OXA-48, KPC, and VIM (n=1), and the VIM gene alone in one strain (n=1). Fourteen isolates were negative for all tested carbapenemase genes (Supplementary Table S1).

Complete genome sequencing revealed that *bla*_NDM-1_ was located on plasmids in 11 isolates and integrated into the chromosome in two isolates, one *E. coli* and one *A. baumannii*. Plasmid sizes ranged from 110,786 bp to 440,281 bp. IncC was the predominant plasmid incompatibility group (n= 6), followed by IncFII (n=3) and IncFIB (n=2) (Supplementary Table S1).

Comparative analysis of the 13 *bla*_NDM-1_-positive isolates revealed substantial diversity in the genetic environments surrounding the resistance genes. A single copy of *bla*_NDM-1_ was identified in each isolate (Figure 1). Despite extensive variation in the surrounding genetic regions, all isolates shared a conserved IS*Aba125*-*bla*_NDM-1_-*ble*_MBL_ core structure. In addition, a downstream *trpF-dsbD*-*CutA*-*groES*-*groEL* module was conserved in eight isolates (10). The diversity of the surrounding genetic structures prompted a detailed investigation of the evolutionary relationships among *bla*_NDM-1_-associated genetic contexts.

**Figure 1:**
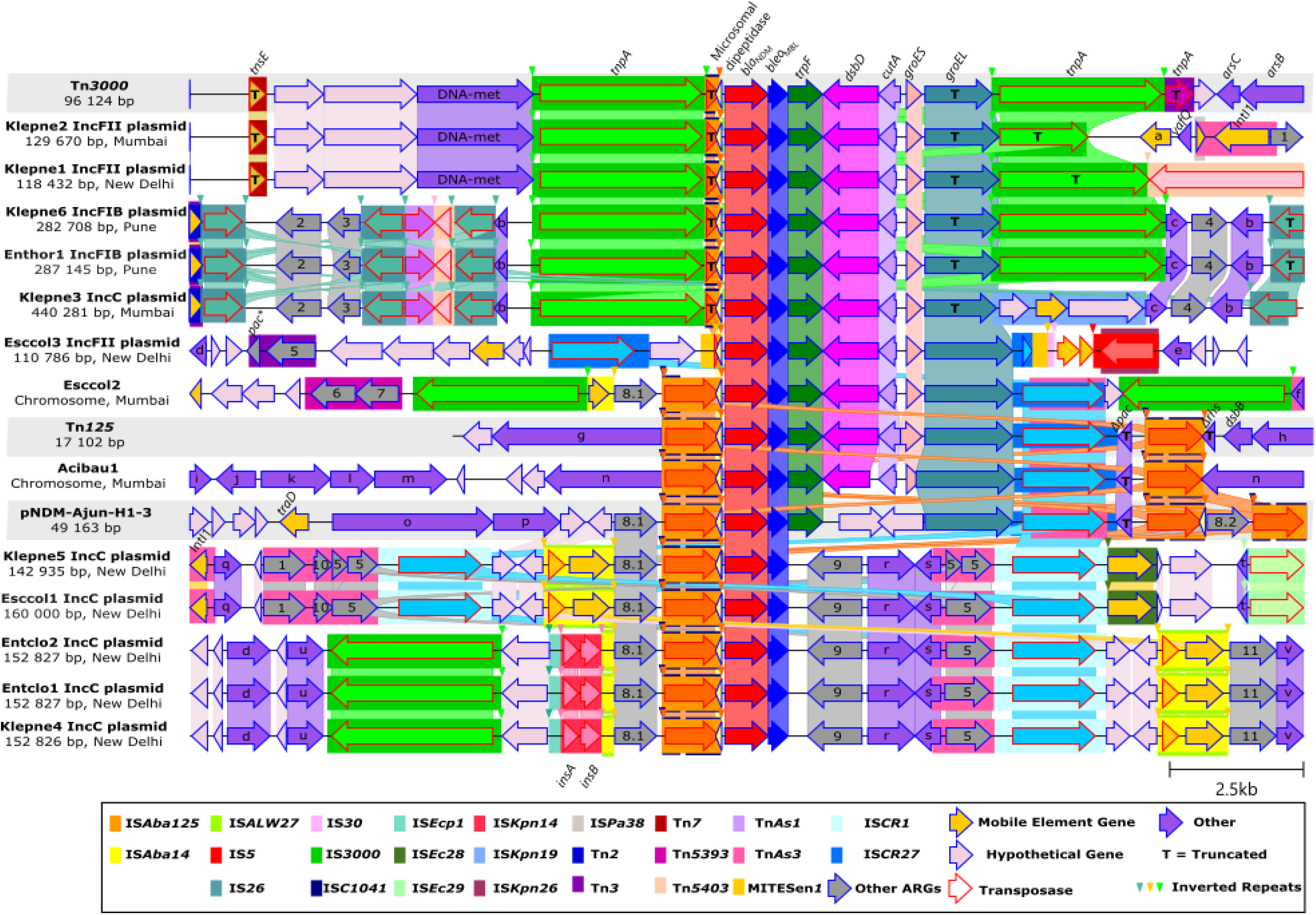
Genetic environment surrounding *bla*_NDM-1_ in 13 Gram-negative bacteria isolated in India in 2007. A Clinker gene cluster comparison of DNA molecules containing *bla*_NDM-1_. Vertical blocks between sequences indicate regions of shared similarity. Genes are drawn to scale and represented by arrows indicating transcriptional orientation. Previously published sequences were highlighted with a gray background. Abbreviations and detailed gene annotations are provided in Supplementary Table S3. Acibau: *Acinetobacter baumannii*; Entclo: *Enterobacter cloacae*; Enthor: *Enterobacter hormaechei*; Esccol: *Escherichia coli*; Klepne: *Klebsiella pneumoniae*.

### Mobile genetic elements associated with *bla*_NDM-1_

Analysis of the 13 newly identified *bla*_NDM-1_ genetic contexts revealed the consistent presence of IS*Aba125* upstream of the resistance gene in all isolates. This element was intact in seven isolates and truncated in six. In contrast, the canonical composite transposon Tn*125*, characterized by two flanking copies of IS*Aba125*, was identified only in the *A. baumannii* isolate (Figure 1). Notably, three copies of IS*Aba125* were previously reported in the oldest *bla*_NDM-1_-carrying sequence described to date (pNDM-Ajun-H1-3) (19).

Tn*3000*-associated structures were identified in nine isolates and exhibited substantial structural diversity. Two isolates (Enthor1 and Klepne6) retained an intact Tn*3000* structure, characterized by transposase genes flanking both ends of the *bla*_NDM-1_-containing region. In contrast, Klepne1 and Klepne2 carried truncated downstream copies of the Tn*3000*-associated transposase gene *tnpA*. In Klepne3, the downstream *tnpA* copy was replaced by IS*Kpn19*. Esccol2 contained two Tn*3000*-associated transposase genes arranged in opposite orientations and linked to distinct flanking regions surrounding the conserved IS*Aba125*-*bla*_NDM-1_-*ble*_MBL_-*trpF-dsbD*-*CutA*-*groES*-*groEL* module. A related organization was observed in Klepne4, Entclo1 and Entclo2, each of which carried a single upstream *tnpA* copy in the opposite orientation.

Among the insertion sequences identified, IS*Aba14*, also known as IS*Aba35*, was particularly notable. Originally described in *A. baumannii* (34), this element was detected in six isolates and was consistently associated with *aphA6,* an aminoglycoside *N*(3)-acetyltransferase resistance gene. In Esccol1, Esccol2 and Klepne5, IS*Aba14* was located upstream of the conserved *bla*_NDM-1_ region. In Entclo1, Entclo2 and Klepne4, two IS*Aba14* copies were present, although the upstream copy was interrupted by an IS*5* element. Notably, IS*Aba14* was also identified in some of the earliest available *bla*_NDM-1_ genetic contexts, including sequences recovered in 2005 and 2006 (22,23,35,36).

ISCR*1* was detected in five isolates. Three carried a single downstream ISCR*1* copy, whereas two contained paired ISCR*1* elements flanking *bla*_NDM-1_ region. ISCR*27* was identified in three isolates and was also present in pNDM-Ajun-H1-3. Esccol3 carried a structure resembling a composite transposon, consisting of a complete upstream ISCR*27* and a truncated downstream copy surrounding *bla*_NDM-1_, whereas Acibau1 and Esccol2 carried only a single downstream ISCR*27* copy.

### Conserved microsomal dipeptidase-like <u>o</u>pen <u>r</u>eading frame (ORF)

A previously overlooked 117-bp ORF encoding a predicted 38-amino-acid protein was identified across newly identified and historical *bla*_NDM-1_ genetic contexts (Figure 1). The same ORF was present in the 13 newly characterized contexts and also in all previously described ancestral *bla*_NDM-1_ structures examined, including pNDM-Ajun-H1-3. BLASTP analyses showed high sequence conservation, with the closest matches corresponding to hypothetical proteins identified in *E. coli* and *A. baumannii* (97.4% and 97.3% identity, respectively). Although annotated as a microsomal dipeptidase-like protein by automated pipelines, the predicted product (MILIWFTIAKIWGEFKHEVRQFQKPYDNQQAEQI SLMV) is substantially shorter than characterized members of this protein family. Sequence variation within the ORF was identified as a single G22A nucleotide substitution, resulting in a conservative valine-to-isoleucine amino acid replacement observed in two sequences (Tn*3000* and pEh1A). Notably, the ORF was consistently oriented opposite to *bla*_NDM-1_ and overlapped with both the IS*Aba125* right inverted repeat (IRR) and the -35 promoter box within the same upstream region of the resistance gene (Figure 2). However, examination of complete IS*Aba125* copies revealed that this ORF is also present in genetic contexts lacking *bla*_NDM-1_, including the duplicated IS*Aba125* regions identified in pNDM-Ajun-H1-3 (Figure 1) and *A. baumannii* plasmid pABF9692 (data not shown). These observations suggest that the ORF is likely a conserved feature of IS*Aba125* rather than a genetic element specifically associated with *bla*_NDM-1_. Nevertheless, its presence in all ancestral and contemporary *bla*_NDM-1_ structures examined suggests that it has been consistently co-mobilized alongside the resistance gene throughout the evolution of Tn*125*-derived elements.

**Figure 2:**
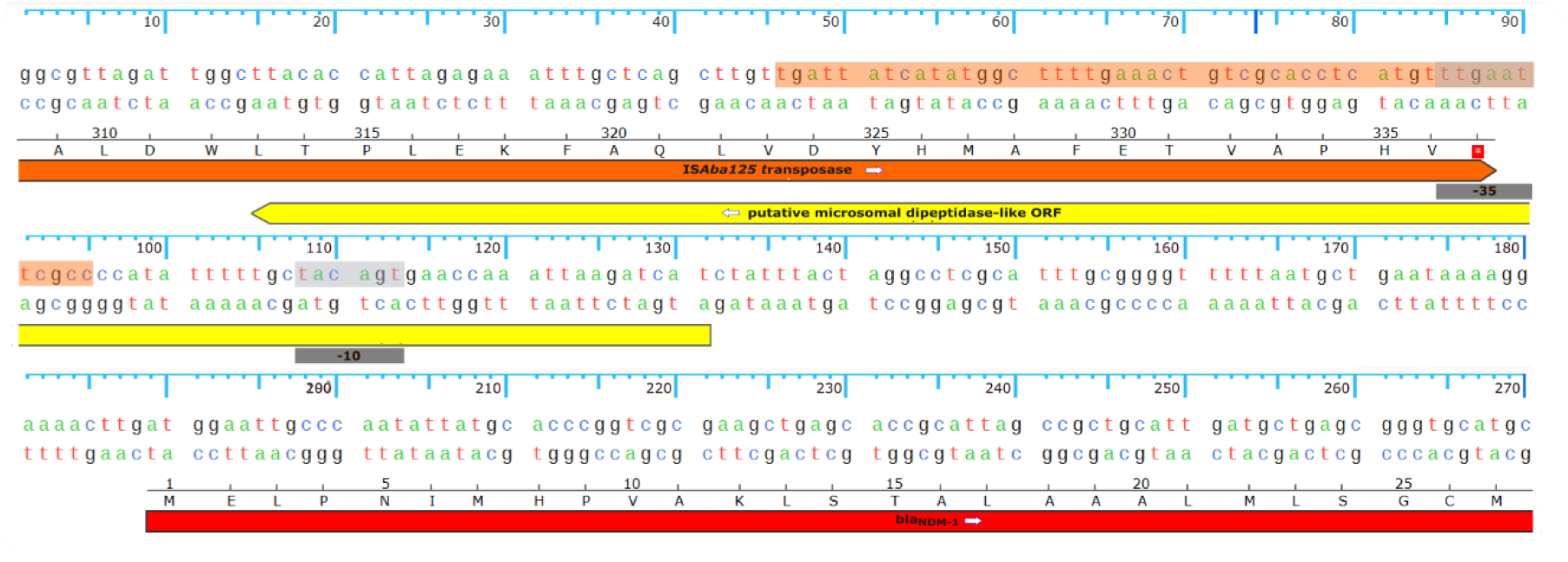
A putative microsomal dipeptidase-like open reading frame encoded within IS*Aba125* overlaps the *bla*_NDM-1_ promoter region. Sequence view generated in Snapgene showing the genetic organization of the region upstream of *bla*_NDM-1_ in the *A. junii* plasmid pNDM-Ajun-H1-3 (NCBI GenBank accession number: GCA_042509155.1). The predicted -35 and -10 promoter boxes are shown in gray, whereas the right inverted repeat (IRR) of IS*Aba125* is highlighted in orange.

### Resistance genes

In addition to *bla*_NDM-1_, numerous acquired resistance genes were identified (Supplementary Table S4), including determinants conferring resistance to β-lactams, aminoglycosides, quinolones, sulphonamides, tetracyclines, trimethoprims, macrolides, phenicols, colistin, fosfomycin, and rifampicin. Extended-spectrum β-lactamases, aminoglycoside-modifying enzymes, and plasmid-mediated quinolone resistance genes were particularly common. Detailed CARD and ResFinder annotations are provided in Supplementary Table S4.

### Antibiograms

The main broad-spectrum resistance determinants to β-lactam antibiotics that are clinically relevant were determined for the thirteen *bla*_NDM-1_-positive isolates. Overall, isolates were resistant to a mean of 81.0% of the antimicrobial agents tested. All isolates exhibited resistance to most β-lactams, including carbapenems, while several displayed intermediate susceptibilities to some of the β-lactams tested. Colistin showed the greatest *in vitro* activity, with nine isolates categorized as susceptible and two as intermediate (Supplementary Table S2).

## Discussion

The dissemination of *bla*_NDM-1_ has extended far beyond clinical settings, with reports on domestic and livestock animals (37,38), as well as environmental reservoirs, including High Arctic soils with no direct human influence (39). Although *bla*_NDM-1_ is now globally distributed, early epidemiological investigations linked many of the first recognized cases to the Indian subcontinent, leading to the hypothesis that this region played a central role in its early emergence and dissemination (40). However, the earliest known occurrence of a *bla*_NDM-1_-positive isolate was identified in Israel in 2004 (19). Together with reports suggesting the presence of early NDM reservoirs in the Balkans and Middle East (5), the geographic origin and initial dissemination pathways of *bla*_NDM-1_ remain unresolved. These observations also highlight how reconstruction of the evolutionary history of *bla*_NDM-1_ may be influenced by geographical sampling bias and the uneven availability of historical bacterial collections. Consistent with the early prominence of NDM-producers in South Asia, *bla*_NDM-1_ was the predominant carbapenemase detected among our archived Gram-negative isolates collected in India in 2007, with 13 of 36 strains harboring the gene. Most *bla*_NDM-1_-positive isolates originated from New Delhi and were recovered from diverse clinical specimens associated with bloodstream, soft tissue infections and pneumonia. While the broad range of specimen types suggests that our collection captures multiple clinical contexts in which NDM-producing bacteria circulated, the predominance of isolates from New Delhi further illustrates the potential impact of sampling bias on inferences regarding the early distribution of *bla*_NDM-1_.

Current evidence supports the origin of *bla*_NDM-1_ within the *Acinetobacter* genus. Tn*125* has been identified on both chromosomes and plasmids of *Acinetobacter* species (21), and is the composite transposon most strongly associated with *bla*_NDM-1_ in this genus. The recent identification of plasmid pNDM-Ajun-H1-3 in a clinical *A. junii* isolate recovered in Israel in 2004 (19) further supports the hypothesis that the ancestral genetic events leading to the emergence of *bla*_NDM-1_ occurred within *Acinetobacter*. Recent phylogenetic analyses have suggested that the most recent common ancestor of the chimeric *bla*_NDM-1_-containing Tn*125* lineage may predate 1990 and potentially extend back to the mid-twentieth century, indicating that the genetic platform responsible for NDM-1 emergence likely evolved decades before its clinical recognition (10). These findings are consistent with a long-standing association between *bla*_NDM-1_, IS*Aba125* and *Acinetobacter*. One proposed evolutionary scenario suggests that *bla*_NDM-1_ originated through an in-frame fusion between a pre-existing metallo-β-lactamase gene and Tn*AphA6*, a composite transposon composed of *aphA6* flanked by two copies of IS*Aba125* (41). The resulting *bla*_NDM-1_-containing Tn*125* created a mobile platform facilitating *bla*_NDM-1_ dissemination.

The spread of *bla*_NDM-1_ involved multiple evolutionary and ecological transitions mediated by diverse mobile genetic elements and non-clonally related bacterial hosts. Among the insertion sequences associated with *bla*_NDM-1_, IS*Aba125* is the most frequently encountered (7,10). Tn*125* appeared to have played a pivotal role during the earliest stages of dissemination, particularly within *Acinetobacter*, where it is typically found flanked by two copies of IS*Aba125* and remains the dominant insertion sequence associated with the ancestral genetic background of *bla*_NDM-1_. Consistent with this model, we identified a complete Tn*125* structure in the *A. baumannii* isolate (Acibau1), whereas six Enterobacterales carried altered forms lacking the downstream IS*Aba125* copy. This observation supports previous reports suggesting progressive degradation of Tn*125* following its transfer into Enterobacterales Similarly, BJ01-like plasmids, which were highly successful vectors of Tn*125* within *Acinetobacter* species, appear to have been supplanted by alternative plasmid backbones better adapted to Enterobacterales (7).

Over time, Tn*3000* appears to have increasingly replaced Tn*125* as a vehicle for *bla*_NDM-1_ mobilization in Enterobacterales (8,22). First described in 2015, Tn*3000* consists of a 11,823-bp sequence bracketed by two copies of IS*3000* and contains a conserved genetic backbone comprising *bla*_NDM-1_-*ble*_MBL_-*trpF-dsbD*-*CutA*-*groES* module and a truncated *groEL* gene. Tn*3000* was originally identified on plasmids recovered from geographically distant regions, including Brazil, Nepal, Morocco, and India, highlighting its contribution to the international dissemination of *bla*_NDM-1_ across diverse plasmid scaffolds (42). Subsequent large-scale genomic analyses demonstrated that Tn*3000*-associated structures are among the most prevalent *bla*_NDM-1_ genetic contexts in Enterobacterales, supporting the view that Tn*3000* emerged as a major platform for the spread of this carbapenemase following the initial dissemination of Tn*125* (10,43). Consistent with this evolutionary scenario, five Enterobacterales isolates carried Tn*3000*, while four additional isolates contained a single IS*3000* element at variable distances upstream of *bla*_NDM-1_. In three of these four isolates, ISCR*1* was located downstream of *bla*_NDM-1_. These hybrid genetic arrangements may represent transitional or degenerated remodeling of *bla*_NDM-1_-associated mobilization platforms. Notably, two isolates contained structures composed of two ISCR*1* copies flanking the resistance locus, while ISCR*27* was identified in three isolates.

Among insertion sequences identified in this study, IS*Aba14* was present in multiple *bla*_NDM-1_ genetic contexts. Although IS*Aba14* has received comparatively little attention in studies investigating *bla*_NDM-1_ genetic environments, its detection in six isolates from our collection and in some of the earliest available *bla*_NDM-1_-bearing sequences suggests that IS*Aba14* formed part of some of the earliest *bla*_NDM-1_-associated genetic platforms. Although Acman *et al.* comprehensively characterized more than 6,000 genetic contexts surrounding *bla*_NDM-1_, their analyses emphasized recurrent structural variants and major mobile genetic elements, potentially overlooking less prevalent insertion sequences associated with its early dissemination such as IS*Aba14* (10). The distribution and evolutionary significance of upstream elements like IS*Aba14* remain largely unexplored. Alternatively, IS*Aba14*-associated structures may have represented a relatively minor evolutionary lineage that was subsequently displaced by more successful mobilization platforms. Together, these observations highlight the diversity of mobile genetic elements associated with *bla*_NDM-1_ and suggest that multiple mobilization pathways have contributed to its dissemination across bacterial species.

Although *bla*_NDM-1_ has occasionally been identified on bacterial chromosomes, the vast majority of carriage occurs on plasmids, which are considered the primary drivers of its dissemination, especially among Enterobacterales (40). Consistent with this role, the plasmid replicons identified in this study were distributed across multiple bacterial species, suggesting extensive lateral gene transfer among taxonomically distinct hosts. Transfer capabilities have previously been reported for pNDM-BJ01-like plasmids recovered from *Acinetobacter* spp. (19,44–46). Together, these observations further highlight the complementary roles of mobile genetic elements and plasmid dissemination in the global spread of *bla*_NDM-1_ (40).

Beyond the mobile genetic elements directly implicated in *bla*_NDM-1_ mobilization, a conserved open reading frame annotated as a putative microsomal dipeptidase was consistently identified adjacent to *bla*_NDM-1_ in all 13 sequences analyzed (Figure 1), as well as in additional publicly available sequences (data not shown). Despite its apparent conservation across diverse *bla*_NDM-1_-associated genetic contexts, this feature has received little attention in previous studies, and when reported, it has generally not been characterized further (47). The current annotation should be interpreted with caution. Although bacterial dipeptidases are typically metallopeptidases involved in peptide metabolism and nutrient acquisition, the predicted protein identified here is only 38 amino acids in length, considerably shorter than previously characterized bacterial peptidases, which generally exceed 450 amino acids. Moreover, it lacks the conserved catalytic motifs associated with known dipeptidase families, suggesting that this sequence may represent a truncated open reading frame, an annotation artefact generated by automated prediction pipelines, or a small protein with an as-yet unknown function. Notably, homologous copies of this ORF were also identified within IS*Aba125* sequences not associated with *bla*_NDM-1_, suggesting that it may constitute an intrinsic feature of IS*Aba125* rather than a genetic element specifically linked to the resistance gene. Nevertheless, its remarkable conservation across IS*Aba125*-containing structures, including all ancestral and contemporary *bla*_NDM-1_ contexts examined here, indicates that it has been inherited during the mobilization and evolution of Tn*125*-derived elements. Whether this conservation reflects selective constraints acting on IS*Aba125*, a regulatory interaction associated with the overlapping promoter region, or an unidentified biological function remains unclear. Future transcriptomic, proteomic, and functional studies will be required to determine whether this small ORF is expressed and to clarify its potential role, if any, in the biology of IS*Aba125* and the maintenance or dissemination of *bla*_NDM-1_-bearing genetic elements. This study provides insight into the diversity of genetic contexts associated with *bla*_NDM-1_ during the early stages of its dissemination among Enterobacterales. Several of the mobile genetic elements identified, including Tn*125*, Tn*3000*, ISCR*27*-associated structures, are consistent with previous reports supporting the robustness of our findings. Nevertheless, several limitations should be considered when interpreting these results. First, the predominance of isolates originating from New Delhi may have introduced geographic sampling bias and may not fully represent the diversity of *bla*_NDM-1_ genetic contexts circulating throughout the Indian subcontinent during this period. Second, although these isolates predate the initial description of *bla*_NDM-1_, they were collected approximately three years after the earliest currently known *bla*_NDM-1_-positive isolate, *Acinetobacter junii* Ajun-H1-3 recovered in Israel in 2004 (20). However, they represent one of the earliest known collections of *bla*_NDM-1_-positive Enterobacterales, contemporaneous with previously reported isolates recovered in India in 2006–2007 for which genomic sequences remain unavailable (23). This study thus provides the earliest available whole-genome sequences of *bla*_NDM-1_-positive Enterobacterales, offering insights into the initial stages of the dissemination of *bla*_NDM-1_. Given the rapid evolution of mobile genetic elements and plasmids, substantial genetic remodeling may already have occurred prior to their recovery. Consequently, some of the genetic structures described here may reflect relatively early diversification events rather than ancestral configurations originally associated with the emergence of *bla*_NDM-1_.

## Conclusion

We provide insight into the early evolutionary history of *bla*_NDM-1_ by characterizing genetic contexts from some of the earliest known Enterobacterales isolates carrying this resistance determinant. Our findings support a model in which *bla*_NDM-1_, initially associated with the ancestral Tn*125* element in *Acinetobacter*, underwent rapid diversification following its introduction into Enterobacterales. In addition to conserved Tn*125* structures, we identified a range of genetic arrangements involving Tn*3000*, IS*Aba14*, ISCR*1* and ISCR*27*, including hybrid configurations that may represent transitional stages during the evolution of *bla*_NDM-1_-associated mobile genetic elements. Together, these observations highlight the remarkable plasticity of the genetic vehicles responsible for the dissemination of this carbapenemase gene.

Given the continuing global spread of *bla*_NDM_-producing pathogens, *bla*_NDM-1_ remains a major public health concern because β-lactams are used as a first-line treatment for severe infections caused by multidrug-resistant Gram-negative bacteria (10). Compounding this issue, NDM-positive pathogens frequently co-harbor additional resistance genes, driving multidrug resistance and contributing to high morbidity and mortality (12,48). The ongoing emergence of novel NDM variants underscores the adaptability of this resistance mechanism.

The genetic events that facilitated the emergence of *bla*_NDM-1_ in Enterobacterales and drove its earliest mobilization nonetheless remain incompletely resolved. Continued sequencing and retrospective analysis of archived collections from diverse geographic regions will be essential to reconstruct these pathways and to identify the ecological and genetic conditions that favored them. Beyond *bla*_NDM-1_, such work would clarify the drivers of resistance gene mobilization more broadly and strengthen our ability to anticipate comparable emergence events.

## Data Summary

**Supplementary Table S1.** Description of 36 strains used in this study with associated epidemiological information.

**Supplementary Table S2.** Antimicrobial profile of *bla*_NDM-1_-positive strains used in this study.

**Supplementary Table S3.** Description of genes and genetic features identified in Figure 1.

**Supplementary Table S4.** Description of antibiotic resistance genes found in *bla*_NDM-1_-positive strains.

## Author statements

### Author contributions

Nancy Allard, Data curation, Formal analysis, Visualization, Supervision, Writing – original draft | Dalila Touati, Formal analysis, Writing – original draft | Nathan Faille, Investigation | Frédéric Grenier, Conceptualization, Data curation, Data acquisition, Data curation, Initial visualization, Formal analysis, Investigation, Methodology, Writing – review & editing | Emy Beauchemin-Lauzon, Data curation, Visualization | Simon Lévesque, Formal analysis, Supervision, Validation, Writing – review & editing | Sébastien Rodrigue, Formal analysis, Validation, Writing – review & editing | Louis-Patrick Haraoui, Conceptualization, Data acquisition, Formal analysis, Funding acquisition, Investigation, Project administration, Resources, Supervision, Validation, Writing – review & editing | All authors reviewed and approved the final manuscript.

## Acknowledgments

The authors would like to thank JMI laboratory for sharing strains and associated metadata.

